# Spatio-temporal variation of skeletal Mg-calcite in Antarctic marine calcifiers in a global change scenario

**DOI:** 10.1101/502765

**Authors:** Blanca Figuerola, Damian B. Gore, Glenn Johnstone, Jonathan S. Stark

## Abstract

Human driven changes such as increases in oceanic CO_2_, global warming and pollution may negatively affect the ability of marine calcifiers to build their skeletons/shells, especially in polar regions. Here we address, for the first time, spatio-temporal variability of skeletal Mg-calcite using bryozoan and serpulid species as models in a recruitment experiment of settlement tiles in East Antarctica. Mineralogies were determined using X-ray diffractometry for 754 specimens belonging to six bryozoan species (four cheilostome and two cyclostome species) and two serpulid species from around Casey Station. All species had calcitic skeletons. Intra- and interspecific variability in wt% MgCO_3_ in calcite among most species contributed to the biggest source of variation overall. Therefore, biological processes seem to be the main factor controlling the skeletal Mg-calcite in these taxa. However, spatial variability found in wt% MgCO_3_ in calcite could also reflect local impacts such as freshwater input and contaminated sediments. The vulnerability of these species to global change is also examined and those species with high-Mg calcite skeletons and low thermal tolerance (e.g. *Beania erecta*) could be particularly sensitive to near-future global ocean chemistry changes.

## Introduction

Increases in oceanic CO_2_, global warming (GW) and pollution will lead to dramatic changes in global ocean chemistry, particularly the carbonate system, in the near future. Some of the expected consequences of the increase in oceanic CO_2_ are a reduction in seawater pH of 0.3–0.5 pH units by 2100 (ocean acidification; OA), a decrease in the carbonate saturation state (Ω) [1] and metal speciation in seawater [2]. In particular, polar regions are acidifying at a faster rate than elsewhere [3]. Temperature increases will also affect the stability of CaCO_3_ [4], although the combined effects of OA and GW remain poorly constrained.

These human driven changes might negatively affect the ability of marine calcifiers to build their calcified skeletons/shells. Marine calcifiers produce a variety of mineralogical forms (polymorphs) including aragonite, calcite and calcite minerals containing a range of magnesium (Mg) content. Organisms with high-Mg calcite structures are predicted to be more vulnerable to OA as the solubility of calcite increases with its Mg-calcite content [5]. The Mg-calcite in echinoderm skeletons, for example, is expected to increase with GW [6], although this increase may be limited and solubility may not increase dramatically at higher ocean temperatures. Organisms at high latitudes may also be more susceptible to contamination (e.g. trace metals and Persistent Organic Pollutants (POPs)) compared to other regions due to their slower metabolism, growth and larval development and consequent slower detoxification processes, and slower colonisation rates [5, 7–10]. Synergistic effects of anthropogenic driven environmental change could mean a significant degradation or even loss of calcareous reef habitats and sediment deposits used as shelter, substrate and food for many benthic communities [5, 11, 12].

Although seawater carbonate saturation state, pH and temperature play an important role in the incorporation of Mg into calcified structures, other environmental (e.g. salinity and Mg/Ca ratio in seawater) and biological factors (e.g. skeletal growth rate) also mediate this process [13–15]. However, the factors that explain spatial and temporal variations in skeletal Mg-calcite, especially in calcifying organisms inhabiting Antarctic waters, remain unclear.

Direct effects of different stressors on particular species of marine calcifiers can be assessed in laboratory experiments. For example, laboratory studies have demonstrated that OA reduces growth and calcification in a wide range of marine calcifiers [16, 17]. However, experiments in the field provide a better understanding of how the effects of environmental change will manifest on calcified organisms living in complex ecosystems [18]. Such studies allow simultaneous exploration of different physical, biological and anthropogenic factors (e.g. depth, species interactions and pollution) which likely interact in their resulting effects on organisms.

In a changing world, ecosystem engineers such as bryozoans and serpulid polychaetes contribute to habitat resilience, especially in Antarctica where these taxa are widespread and diverse [19–21], through their wide roles such as creating reef frameworks that prevent storm damage and enhance biodiversity [22, 23]. Serpulid polychaetes are amongst the most prominent early colonizers in Antarctica but the CaCO_3_ biomineralization of their calcareous tube is relatively unknown [9, 24, 25]. This study investigates the spatio-temporal variability in skeletal Mg-calcite in Antarctic bryozoans and serpulid polychaetes that secrete variable-Mg calcite skeletons [24, 26–28]. Locations around Casey Station in East Antarctica were chosen for this study as they represent excellent opportunities to evaluate skeletal Mg-calcite variations in undisturbed and human-impacted areas in an Antarctic region. Objectives were to: (1) explore spatial variability in skeletal Mg-calcite across locations and water depths; (2) investigate short-term temporal variability (at 3 and 9 y) in skeletal Mg-calcite and (3) test whether or not there would be differences in mineralogy between impacted (sediments contaminated with petroleum hydrocarbons and heavy metals) and non-impacted areas around Casey Station.

## Materials and Methods

### Study area

The study area is located around Casey Station in the Windmill Islands, on the coast of Wilkes Land, East Antarctica (Fig 1). The shallow marine benthic environment is in places muddy sand, gravel, cobbles and boulders [29]. The experiment was conducted at five locations: Brown Bay outer and Brown Bay inner, O’Brien Bay-1 and O’Brien Bay-2 and Shannon Bay. These bays are covered by sea ice (1.2–2 m thick) for most of the year and are ice free for 1–2 months of the year in Brown Bay and 3–4 months in the other bays. Sea ice reduces physical disturbance of the seabed by icebergs and from wave or wind induced turbulence.

**Fig. 1.**
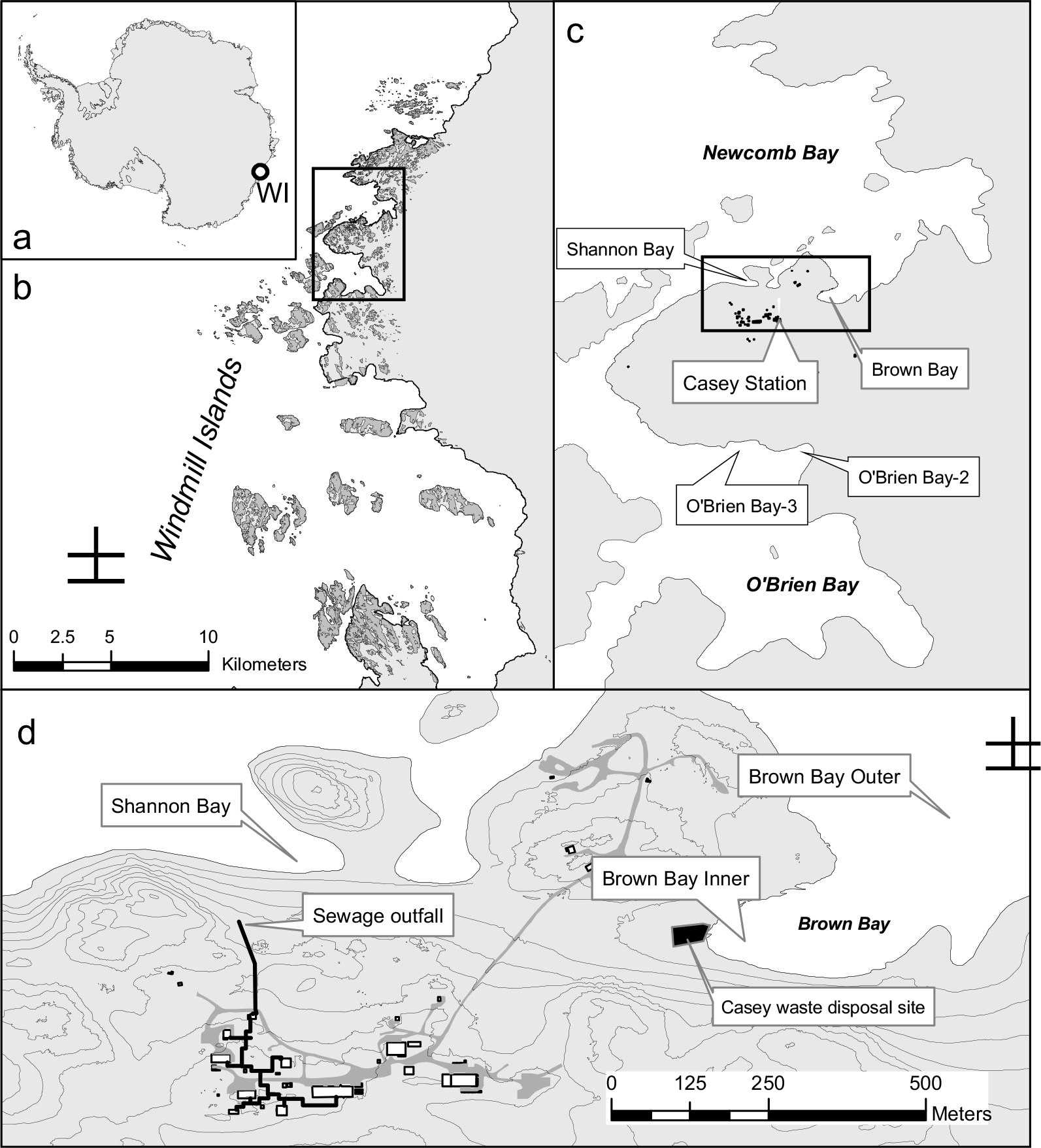
Location of study site: a) Location of Windmill Islands (WI) in East Antarctica; b) the Windmill Islands area with inset c) Newcomb and O’Brien Bays showing the position of the stations with inset d) Casey station, and Shannon and Brown Bays.

Brown Bay is in the southwestern corner of Newcombe Bay. The southern shore is composed of ice cliffs and its western and northern shores are an ice bank at the end of Thala Valley, the location of a former waste disposal site [30]. During summer, a melt stream flows down Thala Valley into Brown Bay. O’Brien Bay is on the southern side of Bailey Peninsula and the shore consists of ice cliffs, 2–30 m high [29]. Shannon Bay is bordered by ice cliffs 2–15 m high and has a wastewater outfall discharging into the bay. Brown Bay and Shannon Bay are adjacent to sources of contamination (~50 m from the waste disposal site and ~100 m from the sewage outfall, respectively) and petroleum hydrocarbons and heavy metals have been detected in sediments [31].

### Sampling design

Two tiles were attached 10 cm apart to the top edge of a trough made from one-half of a 40 cm long, 15 cm diameter stormwater pipe (see the detailed experimental design in [9]). Tiles were deployed at five locations: two human impacted (Brown Bay Inner (BBShall), adjacent to an abandoned waste disposal site, and Shannon Bay (SB), adjacent to a sewage outfall) and three controls (Brown Bay Outer (BBDeep) (500 m from the former waste disposal site), O’Brien Bay-2 (OB2) and O’Brien Bay-3 (OB3)). Tiles were deployed at two depth ranges at each location shallow (6 to 12 m) and deep (15 to 25 m). BBShall had no deep site and BBDeep had no shallow site, due to bathymetry at these locations. At each depth, tiles were deployed at two plots ~20 m apart in groups of 8 tiles (4 troughs ~1 to 2 m apart), with a total of 32 tiles at each location. However, at BBShall and BBDeep, two groups of 8 tiles (in plots of 4 troughs) were placed ~50 m apart to examine small-scale spatial variations. Tiles were deployed between 15 November and 31 December 1997. Tiles were collected after 3 y (2001) and 9 y (2006). One tile was collected from each of two randomly selected pipes at each site/depth at each sampling time (a total of eight per location).

### Collection and identification

Divers collected the tiles which were then preserved in 95% ethanol (Fig 2). Bryozoan colonies were identified to species level in the laboratory using an optical microscope and guide [32]. In subtidal benthic communities, serpulid polychaetes and bryozoans are characteristic early colonists and dominant species until they are outcompeted by other taxa, particularly by sponges and ascidians (Clarke 1996). Six abundant and widely distributed Antarctic bryozoan species selected for this study were the cheilostomes *Arachnopusia decipiens* Hayward & Thorpe, 1988, *Inversiula nutrix* Jullien, 1888, *Beania erecta* Waters, 1904, *Ellisina antarctica* Hastings, 1945, and the cyclostomes *Disporella* cf. and *Idmidronea* cf. Two abundant serpulid polychaete species were also chosen: one species which forms closely coiled tubes (serpulid sp.1) and another species with tusk-shaped tubes (serpulid sp. 2).

**Fig. 2.**
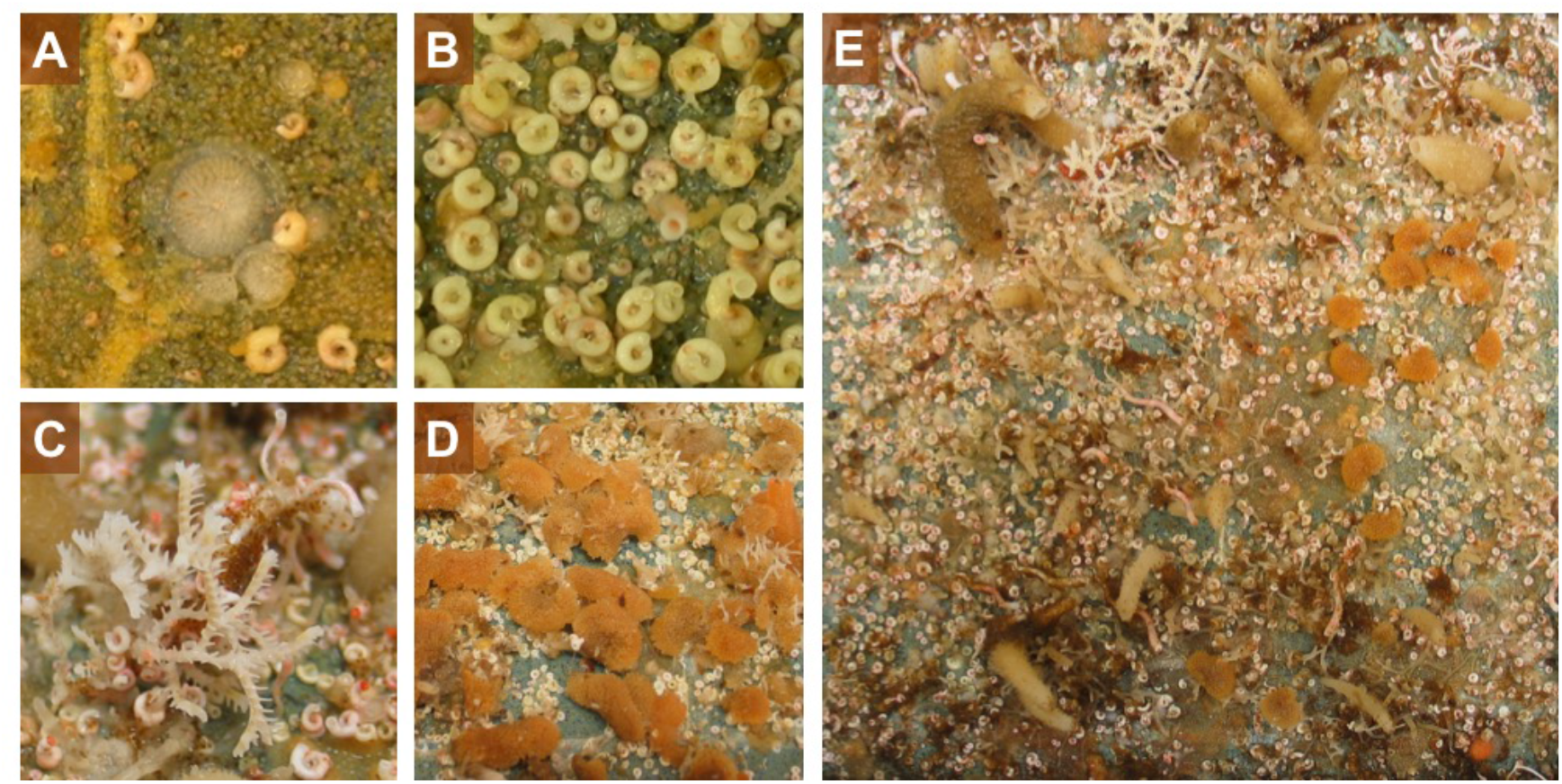
Images of the tiles recovered after 6 (A-B) and 9 (C-E) years. A) A cyclostome colony of *Disporella* cf.; B) serpulid species which form closely coiled tubes (referred here as serpulid sp.1); C) a cyclostome colony of *Idmidronea* cf. and the serpulid species which form tusk-shaped tubes (top right; serpulid sp. 2); D) colonies of the cheilostome *Beania erecta*. E) entire tile colonized by bryozoan colonies, serpulid polychaetes and different species of sponges.

### Mineralogical analyses

Wherever possible, three specimens from each location, each depth, each time and each species were selected for mineralogy analyses. Samples were dissected from 54 tiles with a scalpel blade and a piece (2 mm^2^) from the growing edge of the bryozoan colonies and entire serpulid polychaete tubes were included. After being air-dried, epibionts were removed to avoid mineralogical contamination.

Samples were powdered using an agate mortar and pestle and placed on a Si-crystal low background holder for analysis. X-ray diffractograms were collected from 5 to 90° 2θ with a Panalytical X’Pert Pro MPD diffractometer, using tube conditions of 45 kV, 40 mA, CuK_α_ radiation, X’Celerator detector, Bragg Brentano geometry, and a slew rate of 5° 2θ per minute. Diffractometer performance was constrained by measurement of silver behenate and a single silicon crystal. Limits of detection vary from 0.1 to ~2 wt% depending on the crystallinity of the phase. Identification of minerals, including poorly ordered phases, was conducted using Panalytical’s Highscore Plus software v2.2.4, with ICDD PDF2 and PAN-ICSD databases. Mg-Ca substitution in calcite was quantified via linear interpolation of the angle of the 104 reflection from calcite (CaCO_3_) (0 % Mg-calcite at XX° 2θ) to magnesite (MgCO_3_) (100 % Mg-calcite at YY° 2θ) [33].

Skeletons were divided into low-Mg calcite (LMC; 0–4 wt% MgCO_3_), intermediate-Mg calcite (IMC; 4–8 wt% MgCO_3_) and high-Mg calcite (HMC; >8 wt% MgCO_3_) categories, following a common classification [34].

### Statistical analyses

To test for differences among species, year (age of colony), depth and site, a 4 factor PERMANOVA [35–38] was used. The design included the fixed factors (species, year and depth) and site nested in the combination of species, year and depth. Where significant differences were found, post hoc pairwise PERMANOVA tests were performed. PERMANOVA was done on untransformed skeletal Mg-calcite using a Euclidean distance similarity matrix, which is directly analogous to a univariate ANOVA [39] but is well suited to unbalanced designs, such as is the case in this study (e.g. for depth). The program PRIMER7 (V7.0.13, PRIMER-e, Quest Research Limited) was used for these analyses and the graphical displays were produced using R version 3.5.0 [40]. The package Lattice [41] was called to perform some of the analysis.

## Results

### Intra- and interspecific variation in skeletal Mg-calcite

Mineralogies were determined for 754 specimens belonging to six bryozoan species (four cheilostome and two cyclostome species) and two serpulid polychaete species (Fig 3–6). All species were entirely calcitic.

**Fig. 3.**
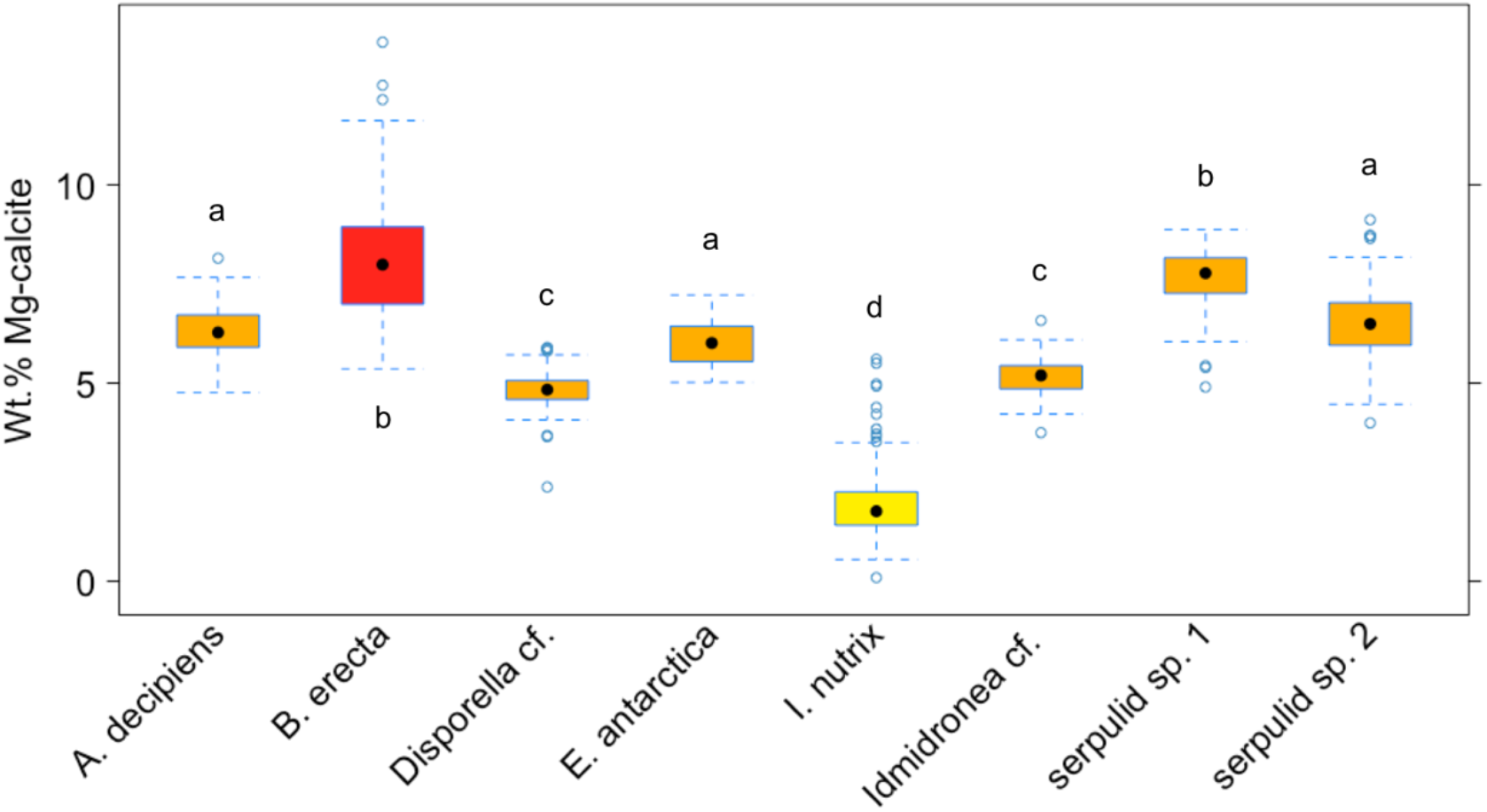
Mean values (±standard error) of wt% MgCO_3_ in calcite in the eight Antarctic species. Boxes depict standard deviation around mean (mid-line); tail indicates range. Colours indicate high-Mg calcite (red), intermediate-Mg calcite (orange) and low-Mg (yellow). Letters indicate results of PERMANOVA tests and show significant differences between species, as indicated by different letters. Species that share the same letter are not significantly different.

**Fig. 4.**
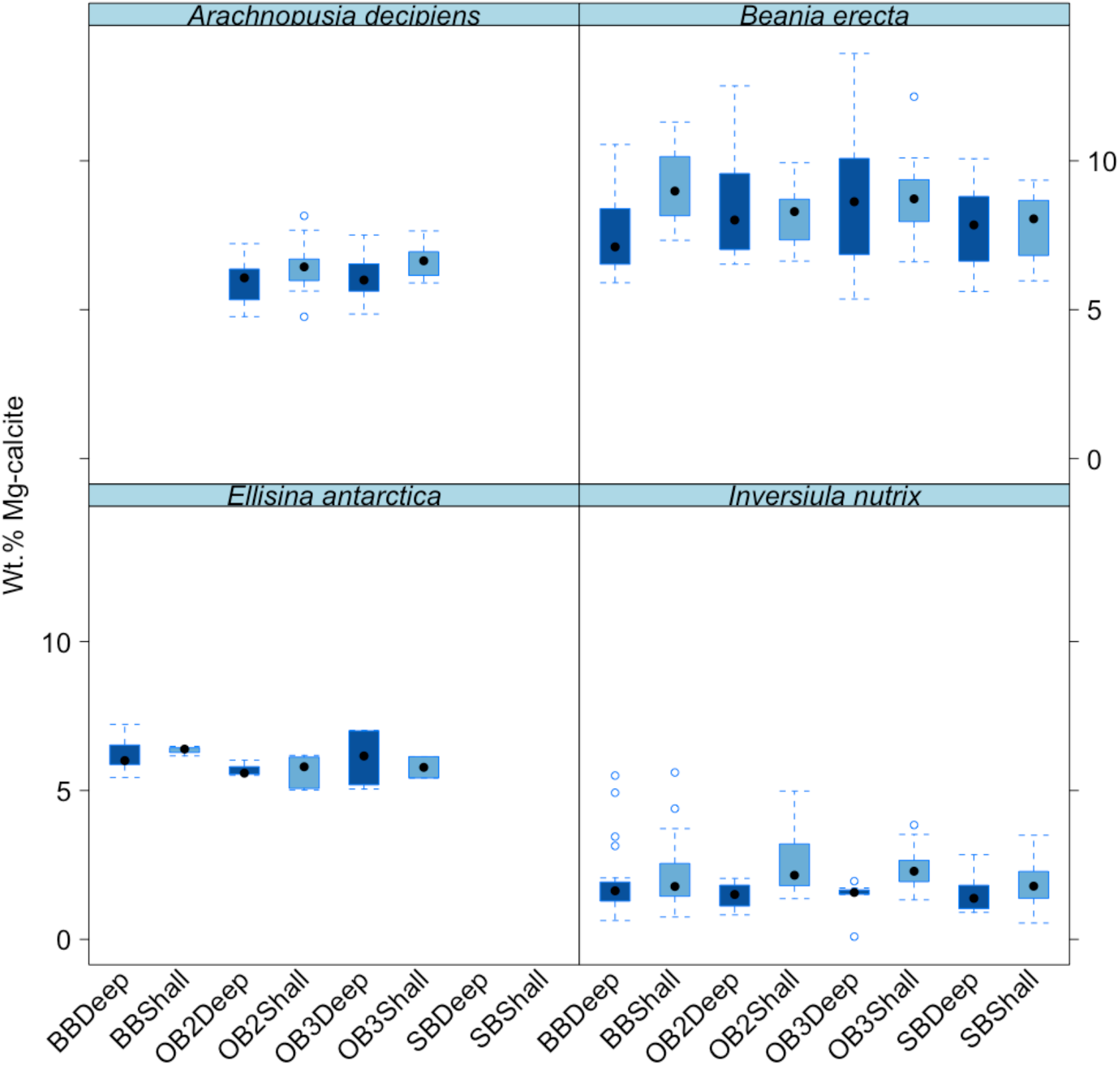
Mean values (±standard error) of wt% MgCO_3_ in calcite in the four Antarctic cheilostome species among different sites and depths. Boxes depict standard deviation around mean (mid-line); tail indicates range. Colours indicate deep and shallow waters. *\*Brown Bay = BB, OB2 = O’Brien Bay −2, OB3 = O’Brien Bay-3, SB = Shannon Bay. Deep = Deep, shallow = Shall*.

**Fig. 5.**
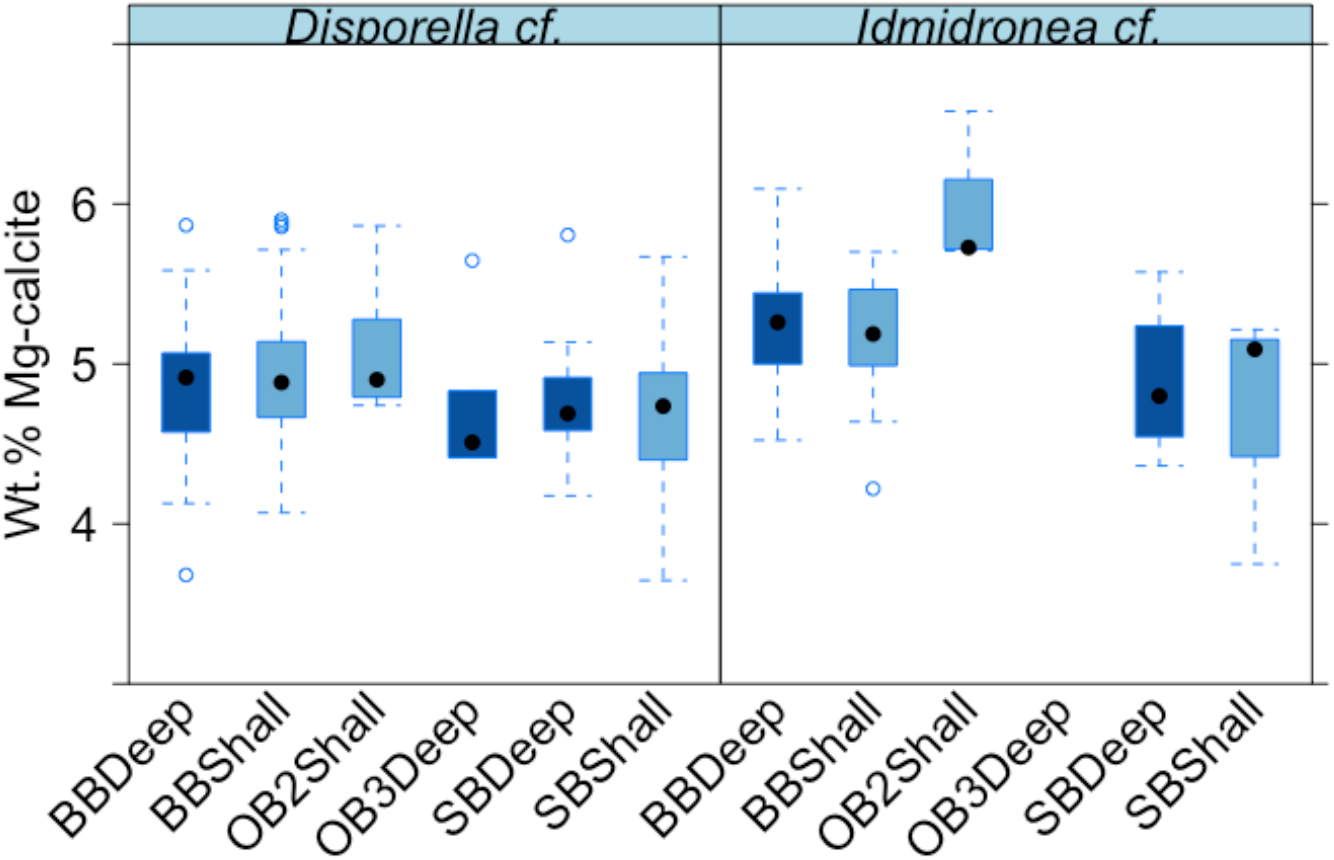
Mean values (±standard error) of wt% MgCO_3_ in calcite in the two Antarctic cyclostome species among different sites and depths. Boxes depict standard deviation around mean (mid-line); tail indicates range. Colours indicate deep and shallow waters. *\*Brown Bay = BB, OB2 = O’Brien Bay −2, OB3 = O’Brien Bay-3, SB = Shannon Bay. Deep = Deep, shallow = Shall*.

**Fig. 6.**
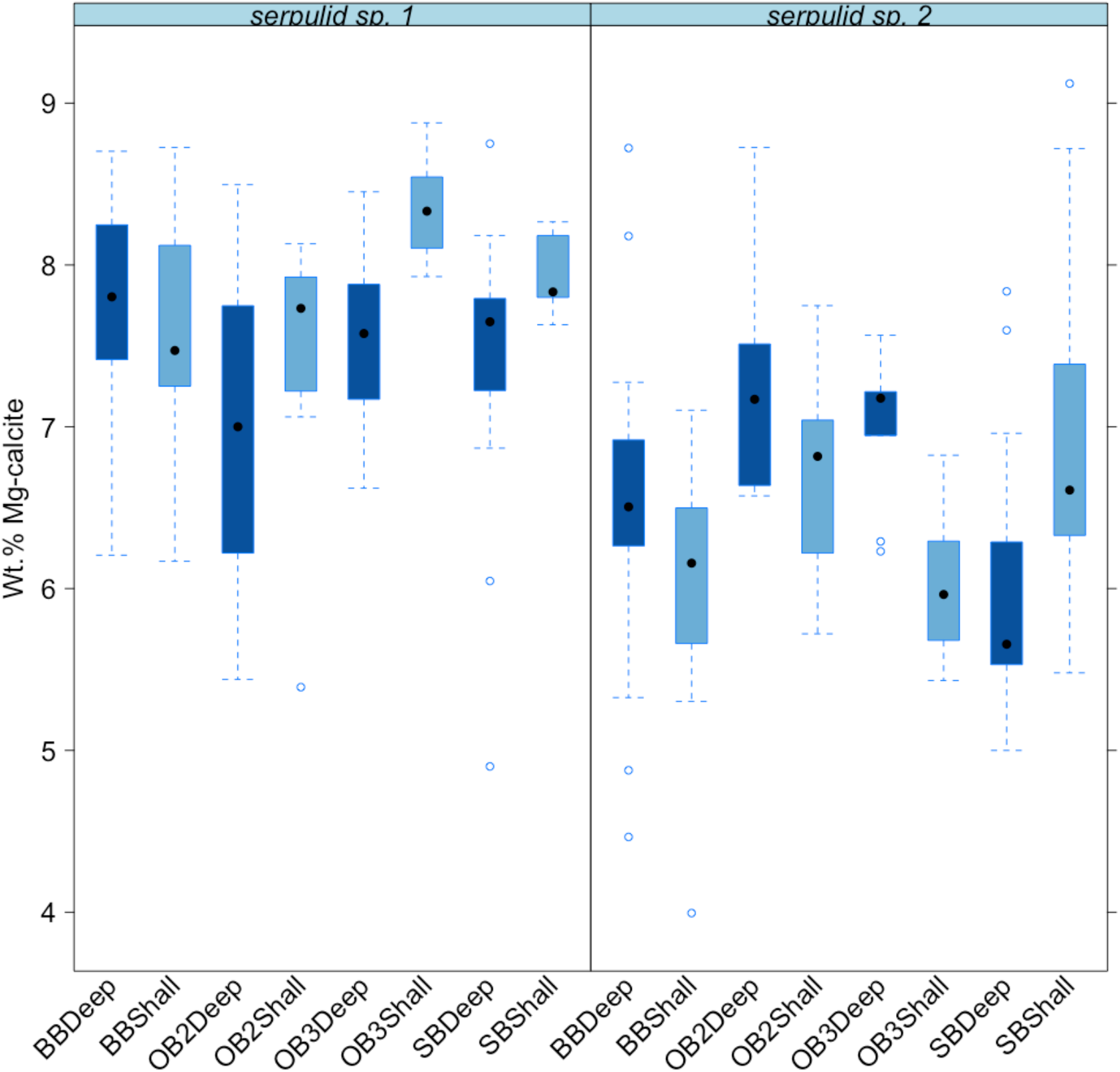
Mean values (±standard error) of wt% MgCO_3_ in calcite in the two Antarctic serpulid species among different sites and depths. Boxes depict standard deviation around mean (mid-line); tail indicates range. Colours indicate deep and shallow waters. *\*Brown Bay = BB, OB2 = O’Brien Bay −2, OB3 = O’Brien Bay-3, SB = Shannon Bay. Deep = Deep, shallow = Shall*.

*Arachnopusia decipiens* (n = 53), *Ellisina antarctica* (n = 28), *Idmidronea* cf. (n = 72) and *Disporella* cf. (n = 111), serpulid sp. 1 (n = 136) and sp. 2 (n = 117) had skeletons of IMC (4–8 wt% MgCO_3_ in calcite) (Fig 3). Of the remaining species, *Beania erecta* (n = 109) consisted of HMC (>8 wt% MgCO_3_ in calcite) and *Inversiula nutrix* (n = 128), of LMC (2–4 wt% MgCO_3_ in calcite).

The pairwise tests for species showed that there is a significant difference in the mean wt% MgCO_3_ in calcite among most species except *Ellisina antarctica* versus serpulid sp. 2 (PERMANOVA pairwise test: *t* = 1.94, *P* = 0.067) and *Arachnopusia decipiens* (*t* = 2.15, *P* = 0.051), serpulid sp. 2 versus *A. decipiens* (*t* = 0.87, *P* = 0.402), serpulid sp. 1 versus *Beania erecta* (*t* = 1.48, *P* = 0.154), *Idmidronea* cf. versus *Disporella* cf. (*t* = 2.23, *P* = 0.051) (Fig 3; Table 1).

**Table 1.** Post hoc pairwise permutational multivariate analysis of variance (PERMANOVA) tests for differences in Mg content between species. Bold type indicates significant difference at p = 0.001, except where *p<0.01.

|  |  | A. | *Disporell* | *Idmidrone* | B. | serpuli | serpuli |
| --- | --- | --- | --- | --- | --- | --- | --- |
|  | *I. nutrix* | decipiens | *a* cf. | *a* cf. | *erecta* | d sp. 1 | d sp. 2 |
| *E. antarctica* | **\*17.25** | 2.15 | **7.70** | **4.12** | **5.33** | **7.71** | 1.94 |
| Serpulid 2 | **24.95** | 0.87 | **9.49** | **6.01** | **6.03** | **6.12** |  |
| Serpulid 1 | **35.76** | **6.76** | **19.02** | **13.34** | 1.48 |  |  |
| *B. erecta* | **27.55** | **5.20** | **14.17** | **10.38** |  |  |  |
| *Idmidronea* cf. | **17.32** | **\*6.03** | 2.23 |  |  |  |  |
| *Disporella* cf. | **18.66** | **10.37** |  |  |  |  |  |
| *A. decipiens* | **21.29** |  |  |  |  |  |  |

### Spatio-temporal analysis in skeletal Mg-calcite

There was a strong significant difference in the mean wt% MgCO_3_ in calcite among species (PERMANOVA pseudo-F = 226.9, p = 0.001) and sites (pseudo-F = 2.04, p = 0.001) (Fig 3–6; Table 2). There was a significant difference among depths (pseudo-F = 5.08, p = 0.029). There were no significant interactions between species, year or depth, indicating differences between species and depth were consistent. There was no effect of year (age of colony).

**Table 2.**
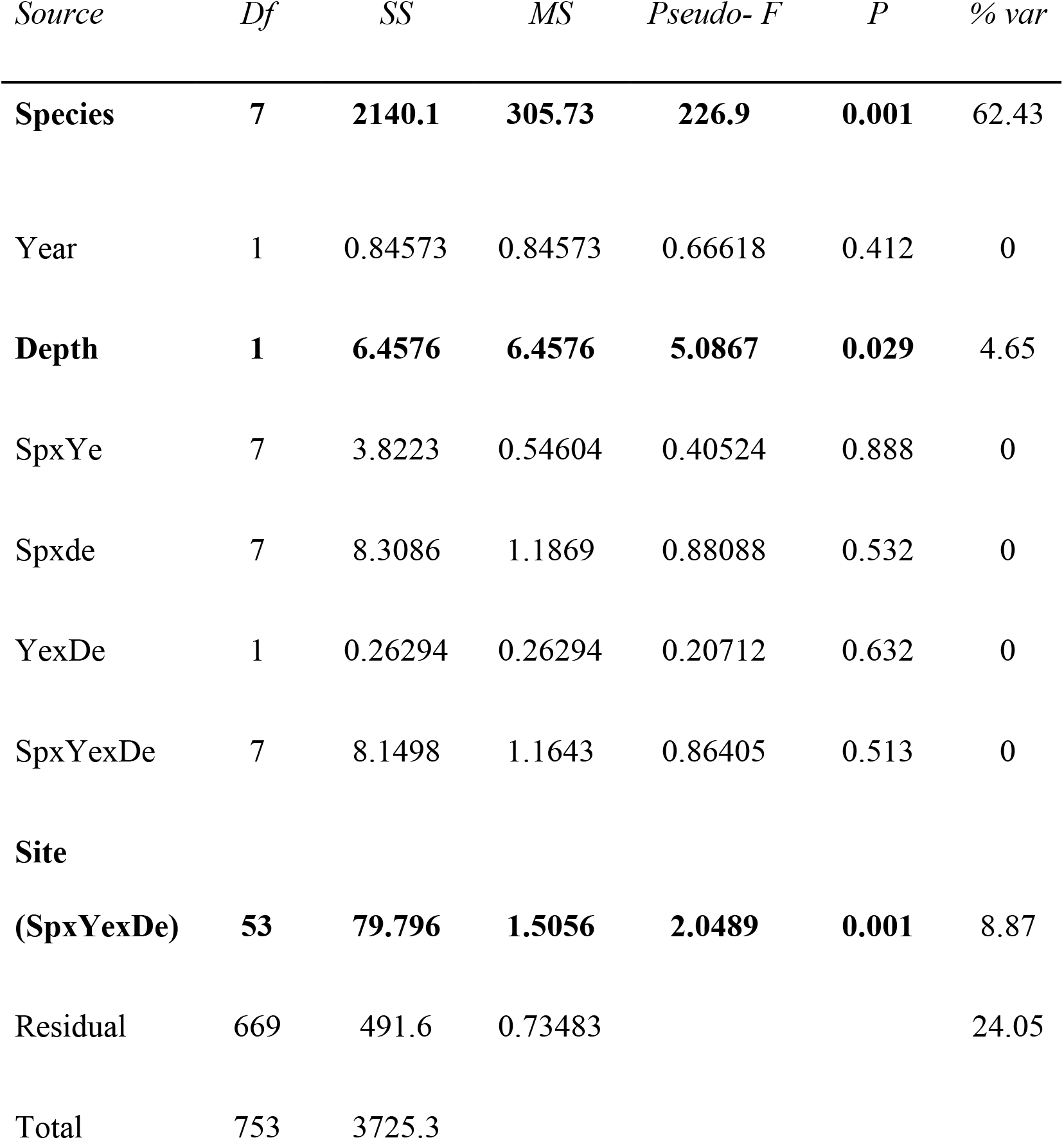
PERMANOVA testing for the effects of species, year and depth on the Mg content in calcite. % var = Estimates of components of variation of Mg content in calcite for each factor in the analysis.

| *Source* | *Df* | *SS* | *MS* | *Pseudo- F* | *P* | *% var* |
| --- | --- | --- | --- | --- | --- | --- |
| **Species** | **7** | **2140.1** | **305.73** | **226.9** | **0.001** | 62.43 |
| Year | 1 | 0.84573 | 0.84573 | 0.66618 | 0.412 | 0 |
| **Depth** | **1** | **6.4576** | **6.4576** | **5.0867** | **0.029** | 4.65 |
| SpxYe | 7 | 3.8223 | 0.54604 | 0.40524 | 0.888 | 0 |
| Spxde | 7 | 8.3086 | 1.1869 | 0.88088 | 0.532 | 0 |
| YexDe | 1 | 0.26294 | 0.26294 | 0.20712 | 0.632 | 0 |
| SpxYexDe | 7 | 8.1498 | 1.1643 | 0.86405 | 0.513 | 0 |
| **Site** |  |  |  |  |  |  |
| **(SpxYexDe)** | **53** | **79.796** | **1.5056** | **2.0489** | **0.001** | 8.87 |
| Residual | 669 | 491.6 | 0.73483 |  |  | 24.05 |
| Total | 753 | 3725.3 |  |  |  |  |

Differences in skeletal Mg-calcite between species contributed to >62% of the total variation observed (Fig 3; Table 2). Variation within species (residual variation among replicates within same depth, year and site) contributed ~24 % to overall variation. Differences among sites (which are confined to comparisons among the same species, depths and years as this was a nested factor in the analysis, i.e. site was nested in species, year and depth) contributed ~9 % of the variance (Table 2, Fig 4–6).

There was a small but significant effect of depth, with differences in skeletal Mg-calcite between different depths contributing ~5 % to the estimated total variation (Fig 4–6).

There was no significant overall effect of human impact on skeletal Mg-calcite concentrations (Table 3) nor any significant effect on any individual species (species x impact interaction, Table 3).

**Table 3.** Results of PERMANOVA analysis testing for the effects of human impact on the Mg content in calcite, with tests for each species (impact x species). % var = Estimates of components of variation of Mg content in calcite.

| *Source* | *df* | *MS* | *Pseudo-F* | *P(perm)* | *% var* |
| --- | --- | --- | --- | --- | --- |
| Impact | 1 | 0.25913 | 0.17276 | 0.69 | 0 |
| Species | 7 | 350.56 | 166.13 | 0.001 | 66.7 |
| Impact x<br>species | 6 | 1.0093 | 0.50947 | 0.769 | 0 |
| Site (Sp. x<br>impact) | 20 | 2.4276 | 3.1551 | 0.001 | 8.4 |
| Residual | 719 | 0.76941 |  |  | 25 |
| Total | 753 |  |  |  |  |

## Discussion

This study evaluates the variability of wt% MgCO_3_ in calcite in the skeletons of a large number of specimens of common bryozoan and serpulid taxa in coastal Antarctic waters, one of the regions expected to be first affected by OA and GW in the near future. To our knowledge, this is the first study addressing the spatio-temporal variability of skeletal Mg-calcite using bryozoan and serpulid polychaete species as models in an experiment of settlement tiles in Antarctica. These two taxonomic groups, and some genera and species studied here (e.g. *Arachnopusia, Beania, E. antarctica*), are common components of hard-substratum sessile assemblages across a wide range of cold latitudes (e.g. South American Region) [42]. These kinds of studies allow regional and latitudinal comparisons of the effects of changes in global ocean chemistry on marine calcifiers. All species had calcitic skeletons, which is consistent with our expectations from previous research on bryozoans and serpulids from high latitudes [28, 43–46]. This trend is possibly due to low temperatures favouring the deposition of calcite over aragonite as the latter is more susceptible to dissolution in cold waters [45].

### Inter- and intraspecific variability in skeletal Mg-calcite

Inter-specific differences in the mean wt% MgCO_3_ in calcite were the largest source of variation overall. Remarkably, there were significant differences in the skeletal Mg-calcite between all bryozoan species except between *E. antarctica* and *A. decipiens* and between the two cyclostome species. There was a wide range of wt% MgCO_3_ in calcite measurements in the bryozoan and serpulid polychaete species studied here, from 0.1–13.1 wt% MgCO_3_ in calcite. These values are not surprising for wt% MgCO_3_ in calcite in bryozoans, which can range from 0 to 14 [43, 44], nor for serpulids, which range from 7 to 15 [24]. Only one species was classified as LMC (*Inversiula nutrix*); five as IMC (*Arachnopusia decipiens, Ellisina antarctica, Disporella* cf., *Idmidronea* cf. and serpulid sp. 1 and sp. 2); and one as HMC (*Beania erecta*). Most cheilostome bryozoans secrete IMC [43, 44], which is further confirmed by this study. To our knowledge, this study also represents the first mineralogical measurements made for the species *A. decipiens*. Our values are within the range of wt% MgCO_3_ in calcite obtained by previous studies in Antarctic cheilostome bryozoans except for *B. erecta*, which has been reported to be IMC [28, 47, 48]. The skeletons of *B. erecta* in our study showed higher mean Mg values even though the range was highly variable (5.3–13.6 wt% MgCO_3_ in calcite). This species exhibits a wide geographical distribution in Antarctica and it could explain the broad range of wt% MgCO_3_ in calcite found here. Other studies have reported ICM and LMC skeletons in *E. antarctica* and *I. nutrix*, respectively, from King George and Adelaide Islands (Antarctic Peninsula)[26, 28, 44]. Significant differences were found in the skeletal Mg-calcite even between cheilostomes of the same suborder Flustrina, in *B. erecta* and *E. antarctica*, although they belong to different families (Beaniidae and Ellisinidae, respectively). Coinciding with our results on the cyclostome genera, *Idmidronea* sp. and *Disporella* sp. produced LMC and HMC in a previous study [43].

The next biggest source of variation was among individuals of the same species. Therefore, biological processes, also known as vital effects [49], seem to be the main factors controlling the skeletal Mg-calcite in these organisms and, in the case of bryozoans, at intra- and interspecific and intracolonial levels [50]. Other studies provide corroborative evidence that the variation in the skeletal Mg-calcite in serpulid polychaete species [51] and Antarctic bryozoans is partially biologically mediated [26, 28].

### Potential environmental influence on skeletal Mg-calcite

Differences among sites (within species) contributed less to variation in skeletal Mg-calcite than among species but was still a significant effect. Several studies have demonstrated that marine calcifiers such as bryozoans, coccoliths, foraminifera and sea stars can respond differently to a range of environmental factors (e.g. water temperature, alkalinity, salinity and Mg/Ca ratio of seawater)[14, 26, 27, 52–54]. The observed values could reflect a relatively stable environment below ice with constant low seawater temperatures as these bays are covered by fast sea-ice almost all the year. Therefore, environmental variables such as seawater temperature could have little influence on the skeletal Mg content in calcite in species inhabiting Antarctic waters, as suggested in previous studies focusing on cold water bryozoans [26, 27]. However, the variation observed could be due to variable temporal local impacts such as freshwater input from the melt stream that flows into Brown Bay during summer. Freshwater not only has different chemistries and transports sediment to the bay, but also affects seawater temperature.

Although depth had the least effect, it was significant, with deeper sites having slightly higher Mg content in calcite. In particular, the significant higher values of Mg content in calcite in deeper waters were only found in the case of serpulid sp. 2 from three control locations (BBDeep, OB2 and OB3). However, other species such as *I. nutrix* and serpulid sp.1 from shallow control locations exhibited higher values than deeper samples. Therefore, there was not any clear depth-related pattern. The lack of influence of depth on mineralogy found is probably due to narrow depth range (6 m) that was sampled.

Although there was no evidence of significant human impact on skeletal Mg-calcite concentrations, further studies covering a broader range area and focusing on other species are needed to evaluate whether or not the Mg content in calcite might be indirectly influenced by contaminated sediments. Heavy metal toxicity may reduce growth rates of marine invertebrates such as bryozoans and worms [55, 56], due to the investment in fitness traits (e.g. growth or reproduction) which are compromised by investment in detoxification mechanisms [57]. Also, organisms with slower growth rates tend to deposit less wt% MgCO_3_ in calcite [5]. Therefore, we could expect lower skeletal Mg content in areas exposed to heavy metals.

### Temporal analyses in skeletal Mg-calcite

We document short term temporal analyses in skeletal Mg-calcite (over a 3 year period) for the first time. Other studies investigated potential temporal variation in skeletal Mg-calcite for greater periods from 30 years to millions of years [28, 44]. In our study, there was no effect of time (age of colony). Similarly, another study evaluating temporal variability in skeletal Mg-calcite in the Antarctic bryozoan species *Cellaria diversa* Livingstone, 1928 and *Antarcticaetos bubeccata* (Rogick, 1955) over the past 30 years did not find significant temporal differences [28]. However, the samples came from greater depth ranges (107–205 and 100–251 m, respectively) and from a different location of Admiralty Bay, South Shetland Islands, north of the Antarctic Peninsula, which lies at 62° 10’ S (versus Casey at 66° 17’ S).

### Vulnerability of the Antarctic target species in a changing world

Human-induced changes could affect survival and calcification of some particular Antarctic marine calcifiers studied here, particularly those with HMC skeletons and low thermal tolerance [5, 11, 58]. Serpulid polychaete tubes are mainly composed of the most vulnerable minerals to OA (aragonite, HMC or a mixture of the two) [24]. In our study, *Beania erecta* and serpulid sp. 1 deposited their CaCO_3_ skeletons containing high proportion of magnesium ions (with mean wt% MgCO_3_ in calcite (± SD) of 8.1 ± 1.52 and 7.7 ± 0.69, respectively) although serpulid sp. 1 was classified here as IMC. The solubility of the skeletons of both species is expected to be greater than that of IMC and LMC or even of aragonite. Although some serpulid polychaetes seem to have the ability to vary their tube mineralogy with seawater chemistry changes, such alterations may result in a deterioration of the tube hardness and elasticity [59]. Consequently, these species are highly likely to be threatened under a scenario of global ocean surface pH reductions of 0.3–0.5 units by the year 2100 [1]. Experimental evidence for this is limited, but the Mediterranean bryozoan *Myriapora truncata* (Pallas, 1766), which has a wt% MgCO_3_ in calcite between 8 and 9.5, has been shown to be vulnerable to ocean acidification at a pH of 7.66, with significant loss of skeleton during a short-term experiment [60].

There is an increase of Mg content in cheilostome bryozoans towards lower latitudes, which is attributed to the warmer seawater [44]. In a recent study, higher values of Mg in two common co-occurring Mediterranean bryozoans *M. truncata* and *Pentapora fascialis* (Pallas, 1766) were also found when the colonies were exposed to high temperatures [61]. Serpulid tubes also show an increase in Mg content with increasing water temperatures [51]. In addition, the interactive effects of multiple environmental changes (e.g. warming, acidification and pollution) have to be taken in account as they are occurring together. The interactive effects of temperature and CO_2_ concentrations on the widely distributed cheilostome bryozoan *Jellyella tuberculata* (Bosc, 1802), including from subAntarctic regions [62], was investigated under controlled conditions [63]. Colonies exposed to reduced pH and increased temperature showed dissolution of their zooids, possibly due to an increase of skeletal Mg content at high temperatures which made the skeletons more susceptible to dissolution under high CO_2_ [63]. Therefore, organisms living in Antarctic coastal areas that have experienced significant warming in recent years (e.g. Western Antarctic Peninsula) could become more vulnerable to environmental changes if their Mg content increases with temperature [64], however it is not known whether this is occurring.

Such changes could lead to shifts in community composition of calcareous marine organisms in future oceans, especially in polar regions [11]. Species with HMC, such as *B. erecta* and serpulid sp. 1, may be particularly vulnerable to the combined effects of OA and GW, although variation in species-specific responses may also be anticipated. Potential subsequent decreases of some species like *B. erecta*, which is widely distributed and common throughout Antarctica and has colonies which comprise dense mats and may cover large areas of substratum, could indirectly lead to negative impacts to other organisms which use their colonies as substrate, food and shelter [32]. This could be expected in colonies living in shallow waters of some Antarctic regions more exposed to increased temperatures such as the Antarctic Peninsula. However, *B. erecta*, has a depth range is from 0 to >1500 m [62], and cooler deep waters may form refugia for this and other species. Also, *B. erecta* is a competitive dominant species, covering different bryozoan species like *Inversiula nutrix* studied here and other sessile organisms [65]. Its decrease could favour other less competitive species such as *I. nutrix*, which may be more resilient to acidification due to lower Mg content of its skeleton and, consequently, replacing the ecological niche occupied by *B. erecta*.

Calcifying marine invertebrates are important components of marine ecosystems and it is clear OA will have a range of effects at the species level, potentially leading to wider ecosystem level effects. Given the range and variation in mineralogy of important calcifying species as observed in this study, some species are likely to be more heavily affected than others. Further work is required to better understand the potential impacts of OA on marine benthic communities, particularly in areas thought to be vulnerable to OA such as Antarctica shallow waters.

## Acknowledgements

We thank dive teams at Casey Station for field assistance. We thank Dr. P. D. Taylor for his constructive comments on our paper. The research visit of B. Figuerola to the Australian Antarctic Division and Macquarie University received support from a Council of Managers of National Antarctic Programs (COMNAP) Research Fellowship. This project was supported financially by the Australian Antarctic Division grant AAS 4127.

